# Structural proteomics reveals a coagulation–complement accessibility signature of macrovascular invasion in hepatocellular carcinoma

**DOI:** 10.64898/2026.08.12.744566

**Authors:** Ahrum Son, Moon Haeng Hur, Eun Ju Cho, Jaeho Ji, Eunjeong Han, Yunho Choi, Jumin Park, Heejin Lee, Sujin Park, Su Jong Yu, Hyunsoo Kim

**Author notes:** Corresponding authors **Hyunsoo Kim** — Department of Convergent Bioscience and Informatics, Chungnam National University, Daejeon 34134, Republic of Korea.; **Su Jong Yu** — Department of Internal Medicine and Liver Research Institute, Seoul National University College of Medicine, Seoul 03080, Republic of Korea. These authors contributed equally to this work (Ahrum Son, Moon Haeng Hur).

## Abstract

Macrovascular invasion (MVI) and extrahepatic spread (EHS) define the most aggressive, treatment-refractory hepatocellular carcinoma (HCC), yet blood-based markers that report the underlying protein-network biology are lacking. Conventional proteomics measures protein abundance but not the conformational and protein–protein-interaction (PPI) states that govern function. We applied covalent proteome painting (CPP)—a dimethylation-based accessibility assay that reads out binding-site openness—to matched tumor and serum, reasoning that intravascular tumor dissemination remodels plasma protein complexes in a manner detectable as changes in accessibility.

Eight treatment-native HCC patients were profiled by CPP using matched FFPE tumor and top-14– depleted serum on a Q Exactive Orbitrap HF. The 85 tumor–serum common proteins defined an 81-protein targeted panel, validated by multiple-reaction-monitoring (MRM) mass spectrometry with heavy stable-isotope-standard peptides (296 peptides; 3,717 light/heavy transition pairs) in 22 FFPE tumors and 22 matched sera. Accessibility was the light/heavy ratio (high, open; low, closed). We assessed differential accessibility, serum–tissue translatability, pathway enrichment, and biomarker/survival performance.

Aggressive disease showed broadly decreased protein accessibility. MVI-associated changes were directionally concordant between tumor and serum (Spearman ρ=0.21; 59% concordant), driven by coagulation and complement proteins (FGG, CTSD, LBP, C4BPA); the EHS axis did not translate. Decreased-accessibility proteins were enriched for complement–coagulation cascades and IGF/IGFBP transport. A six-protein serum accessibility signature discriminated MVI (leave-one-out cross-validated AUC 0.80; best single markers ceruloplasmin 0.83 and haemoglobin-α 0.77), and MVI status trended with shorter overall survival (log-rank p=0.06).

Accessibility-based serum proteomics captures MVI-associated protein-complex remodeling that abundance assays miss, nominating a coagulation/complement-anchored serum signature for vascular-invasive HCC that warrants prospective validation.

## Introduction

Hepatocellular carcinoma (HCC) is among the most lethal malignancies worldwide and remains a leading cause of cancer-related death, with incidence and mortality still rising across many regions^1,2^. Although curative options exist for early-stage disease, prognosis deteriorates sharply once the tumor acquires vascular-invasive or metastatic behavior^3,4^. Macrovascular invasion (MVI)—encompassing portal- and hepatic-vein tumor thrombosis—and extrahepatic spread (EHS) are the principal hallmarks of this aggressive phenotype and are decisive for staging and treatment allocation: within the Barcelona Clinic Liver Cancer framework both denote advanced (stage C) disease for which systemic therapy is recommended, and each is associated with markedly shortened survival, frequently on the order of a few months in untreated series^5-7^. Critically, MVI and EHS are ascertained from cross-sectional imaging or surgical pathology rather than from a blood test, and neither directly reports on the molecular processes that drive intravascular tumor dissemination. A minimally invasive readout that captures the biology of vascular invasion—rather than merely its anatomical consequences—would therefore address a substantial and unmet clinical need.

Blood-based proteomics is an attractive route to non-invasive tumor characterization, yet conventional workflows quantify protein abundance^8,9^. However, much of the biology relevant to metastatic spread is encoded not in how much of a protein is present but in its interaction state—which molecular surfaces are engaged and which remain solvent-exposed. Intravascular tumor cells actively co-opt the haemostatic system: they activate platelets, engage the coagulation cascade, and fix complement, assembling tumor-cell–platelet aggregates that shield circulating tumor cells from immune clearance and shear stress, promote their arrest at the endothelium, and support colonization of distant sites^10-12^. These processes manifest predominantly as the formation and remodeling of protein complexes in plasma—a regulatory layer to which abundance-based assays are, by construction, structurally blind.

Structural, or interaction-state, proteomics has emerged to interrogate precisely this blind spot. Approaches such as limited proteolysis–coupled mass spectrometry and thermal-proteome profiling resolve proteome-wide conformational and binding changes as altered protease susceptibility or thermal stability^13-15^, yielding information that is orthogonal and complementary to abundance and to conventional interactomics^16^. Covalent proteome painting (CPP) provides a particularly direct readout of accessibility: reductive dimethylation covalently tags solvent-accessible lysine ε-amines and protein N-termini, whereas residues buried at protein–protein interfaces or within folded complexes are protected and label less efficiently ^17-19^. The resulting labeling intensity is therefore a quantitative measure of local accessibility—high signal marks an open, solvent-exposed binding site, whereas low signal marks a closed site engaged in complex formation. Applied across the proteome and, as demonstrated here, transferred to a reproducible targeted assay, CPP converts protein-complex biology into a measurable serum analyte.

We hypothesized that the aggressive HCC phenotype—particularly the combination of MVI and EHS— drives intravascular tumor-cell penetration that activates platelet, coagulation, and complement systems, increasing protein-complex formation in plasma and thereby decreasing the CPP accessibility of the participating proteins. Aminopeptidase N (ANPEP/CD13), a moonlighting ectoenzyme shed from activated platelet membranes into plasma that participates in angiogenic and proteolytic complexes, provided a concrete mechanistic candidate^20,21^, as did the extensive crosstalk between the complement and coagulation systems that accompanies malignancy^22,23^. To test whether such structural changes are (i) detectable, (ii) shared between tumor and serum, and (iii) informative for vascular-invasion phenotyping, we designed a two-stage study: unbiased CPP discovery across a balanced 2×2 factorial of MVI and EHS, followed by targeted MRM validation of the tumor–serum common proteome in an expanded, sample-count-harmonized cohort.

## Results

### A two-stage covalent-proteome-painting workflow links the tumor and serum structural proteomes of aggressive HCC

To disentangle the contributions of macrovascular invasion and extrahepatic spread, we selected eight treatment-naïve HCC patients in a balanced 2×2 factorial design (MVI± × EHS±, two patients per group) and profiled matched FFPE tumor and pre-treatment serum by CPP (**Figure 1**). Because a small number of highly abundant plasma proteins dominate the serum signal, serum was first depleted of the 14 most-abundant plasma proteins to expose the lower-abundance interactome, and all samples were profiled on a mass spectrometer, identifying 644 tumor and 288 serum proteins. Their intersection—85 proteins detected in both compartments—defined a targeted panel that, after removal of immunoglobulin and cytoskeletal contaminants, comprised 81 proteins suitable for quantitative validation.

**Figure 1.**
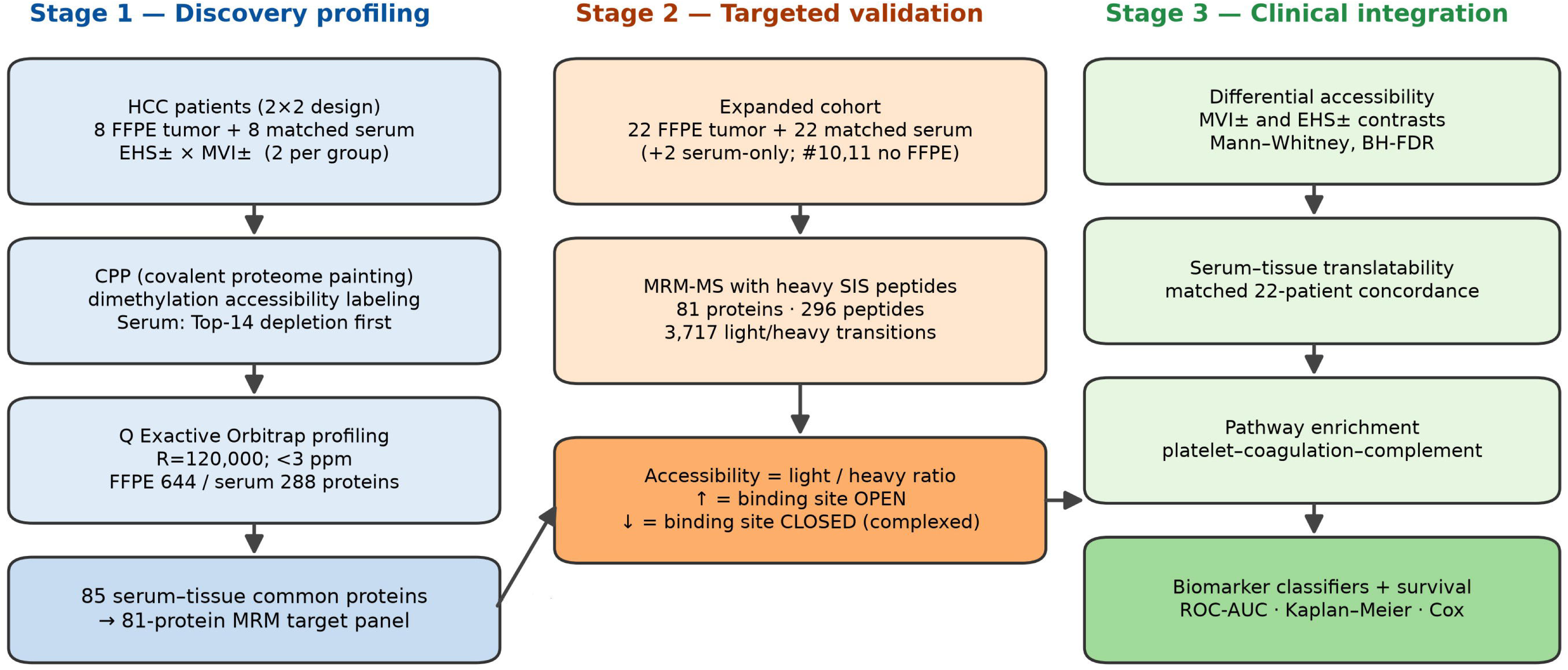
Two-stage covalent-proteome-painting (CPP) structural-proteomics workflow for aggressive hepatocellular carcinoma. Schematic of the study design. *Stage 1 (discovery):* eight treatment-naïve HCC patients were selected in a balanced 2×2 factorial design by macrovascular invasion (MVI) and extrahepatic spread (EHS), two patients per group, and profiled in matched tumor (FFPE) and serum by CPP dimethylation-based accessibility labeling, with serum first depleted of the 14 most-abundant plasma proteins. Global profiling on a Q Exactive hybrid quadrupole-Orbitrap (resolution 120,000; mass accuracy <3 ppm) identified 644 tumor and 288 serum proteins, whose intersection (85 common proteins) defined an 81-protein targeted panel. *Stage 2 (targeted validation):* an expanded cohort of 22 FFPE tumors and 22 matched sera (plus 2 serum-only cases; patients 10 and 11 lacked evaluable FFPE) was analyzed by multiple-reaction-monitoring (MRM) MS with heavy stable-isotope-standard (SIS) peptides—81 proteins, 296 peptides, 3,717 light/heavy transition pairs. Accessibility was quantified as the endogenous-light to SIS-heavy ratio; a higher ratio denotes a more open (accessible) binding site and a lower ratio a more closed (complex-bound) site. *Stage 3 (clinical integration):* differential-accessibility testing across the MVI and EHS axes, serum–tissue translatability, pathway enrichment, and biomarker/survival analysis. The working model posits that intravascular tumor dissemination in EHS(+)/MVI(+) disease activates platelet–coagulation– complement cascades, increasing protein-complex formation and thereby decreasing CPP accessibility (e.g., ANPEP shed from activated platelets).

The validation cohort comprised 22 FFPE tumors and 22 matched sera. Because two patients (cases 10 and 11) lacked evaluable FFPE material, all cross-compartment analyses were harmonized to the same 22 individuals, with two additional serum samples retained only for serum-level estimates (24 sera total). Each protein was quantified by multiple reaction monitoring-mass spectrometry (MRM-MS) with heavy stable-isotope-standard (SIS) peptides (296 peptides; 3,717 transitions), and accessibility was computed as the ratio of endogenous-light to SIS-heavy signal—a high ratio denoting an open, accessible site and a low ratio a closed, complex-engaged site. The assay was both reproducible and complete: all 81 proteins were quantified in every sample, transition-level coefficients of variation were low, and heavy/light transition pairs were tightly correlated (**Figure S1**), establishing a robust quantitative foundation for the differential analyses that follow.

### An advanced, HBV-predominant validation cohort with balanced MVI and EHS representation

The 22-patient validation cohort had uniformly advanced disease (median overall survival 8.0 months; 8 deaths), was predominantly HBV-related (77%), and was largely BCLC stage C (**Table 1**). Macrovascular invasion was present in 16 patients and extrahepatic spread in 9, providing adequate representation of both axes. Stratification by EHS revealed the expected clustering of aggressive features—EHS-positive tumors were more often multifocal (p=0.004) and arose in younger, uniformly male patients—whereas most laboratory and tumor-burden variables did not differ significantly between MVI or EHS strata. The absence of systematic differences in gross tumor burden indicates that the accessibility differences described below are unlikely to be mere proxies for tumor mass, and instead reflect qualitative remodeling of the circulating proteome.

**Table 1.** Clinical and demographic characteristics of the validation cohort. Summary of the 22-patient targeted-validation cohort. Values are counts (percentages) or medians unless otherwise indicated. Full characteristics stratified by MVI and EHS status, including all laboratory and tumor-burden variables.

| Variable | Overall (n=22) | MVI(-) (n=6) | MVI(+) (n=16) | MVI<br>p | EHS(-) (n=13) | EHS(+) (n=9) | EHS<br>p |
| --- | --- | --- | --- | --- | --- | --- | --- |
| Age, years | 58.0 (52.5–63.0) | 64.0 (58.5–74.0) | 55.5 (50.8–62.2) | 0.0970 | 61.0 (56.0–67.0) | 52.0 (50.0–60.0) | 0.061 |
| Male sex, n (%) | 17 (77) | 4 (67) | 13 (81) | 0.5850 | 8 (62) | 9 (100) | 0.054 |
| HBV etiology, n (%) | 17 (77) | 3 (50) | 14 (88) | 0.1000 | 9 (69) | 8 (89) | 0.360 |
| Diabetes, n (%) | 4 (18) | 2 (33) | 2 (12) | 0.2920 | 2 (15) | 2 (22) | 1.000 |
| Hypertension, n (%) | 9 (41) | 2 (33) | 7 (44) | 1.0000 | 7 (54) | 2 (22) | 0.203 |
| Maximum tumor size, cm | 12.2 (8.2–14.9) | 13.6 (11.3–14.8) | 10.9 (8.0–14.6) | 0.5300 | 14.0 (9.6–15.0) | 9.8 (8.0–12.7) | 0.141 |
| Tumor number | 3.0 (1.0–4.0) | 4.0 (4.0–4.0) | 1.0 (1.0–4.0) | 0.1310 | 1.0 (1.0–2.0) | 4.0 (4.0–4.0) | 0.004 |
| Child-Pugh score | 6.0 (5.0–6.0) | 5.5 (5.0–6.0) | 6.0 (5.0–6.2) | 0.7220 | 6.0 (5.0–7.0) | 5.0 (5.0–6.0) | 0.316 |
| Platelets, ×10<sup>3</sup>/μL | 187.0 (129.2–274.8) | 234.5 (187.0–339.0) | 141.5 (116.2–214.8) | 0.1310 | 199.0 (137.0–282.0) | 187.0 (127.0–192.0) | 0.738 |
| Albumin, g/dL | 3.5 (3.2–4.0) | 3.8 (3.5–4.3) | 3.5 (3.0–4.0) | 0.1380 | 3.4 (3.0–4.0) | 3.6 (3.5–4.0) | 0.283 |
| Total bilirubin, mg/dL | 0.9 (0.6–1.4) | 1.1 (0.8–1.6) | 0.9 (0.6–1.3) | 0.3750 | 1.3 (0.6–1.7) | 0.9 (0.5–1.1) | 0.170 |
| AST, IU/L | 134.0 (80.2–202.2) | 134.5 (107.5–190.0) | 127.0 (60.0–197.8) | 0.5900 | 149.0 (110.0–209.0) | 76.0 (50.0–145.0) | 0.045 |
| ALT, IU/L | 61.0 (32.2–110.2) | 72.5 (46.8–79.5) | 47.0 (30.8–140.2) | 0.9140 | 81.0 (42.0–135.0) | 36.0 (27.0–70.0) | 0.142 |
| AFP, ng/mL | 3766.5 (152.6–18602.5) | 6650.9 (74.5–17417.5) | 3766.5 (203.1–20640.0) | 0.8580 | 4470.0 (48.7–28890.0) | 3063.0 (155.4–17890.0) | 0.841 |
| PIVKA-II, mAU/mL | 9306.0 (1656.0–26616.0) | 4887.0 (1879.2–54408.2) | 14488.0 (2682.5–24338.0) | 0.8460 | 19670.0 (3709.0–26616.0) | 4887.0 (243.2–23916.8) | 0.277 |
| EHS-positive, n (%) | 9 (41) | 3 (50) | 6 (38) | 0.6550 | 0 (0) | 9 (100) | — |
| MVI-positive, n (%) | 16 (73) | 0 (0) | 16 (100) | — | 10 (77) | 6 (67) | 0.655 |
| PFS, months | 4.7 (2.2–7.8) | 3.9 (2.4–8.4) | 5.0 (2.1–7.4) | 0.9380 | 5.2 (2.2–7.2) | 3.0 (2.2–9.3) | 0.859 |
| OS, months | 8.0 (4.7–10.8) | 9.2 (5.5–19.7) | 7.8 (4.3–10.5) | 0.5330 | 9.9 (5.4–11.3) | 6.3 (3.7–9.3) | 0.425 |
| Death, n (%) | 8 (36) | 1 (17) | 7 (44) | 0.3510 | 5 (38) | 3 (33) | 1.000 |

*BCLC, Barcelona Clinic Liver Cancer; EHS, extrahepatic spread; HBV, hepatitis B virus; MVI, macrovascular invasion.*

### Aggressive HCC is characterized by a coordinated loss of protein-binding-site accessibility

In the discovery data, the most aggressive group (G1, EHS+/MVI+) exhibited predominantly decreased accessibility relative to the double-negative control group (G4, EHS−/MVI−) across the common proteome (**Figure 2a, c**). Proteins losing accessibility in aggressive disease included carboxylesterase-1, haemoglobin subunits, cathepsin D, and keratins—consistent with sequestration of these species into larger, less solvent-exposed complexes—whereas a smaller set, including apolipoproteins and transferrin receptor, gained accessibility. At the individual-protein level the discovery serum and tissue measurements were only weakly correlated (ρ≈0.05; **Figure 2b**), as expected for n=2 per group; discovery therefore served to nominate candidate proteins and to define the direction of the dominant effect, with quantitative and statistical claims deliberately reserved for the larger validation cohort.

**Figure 2.**
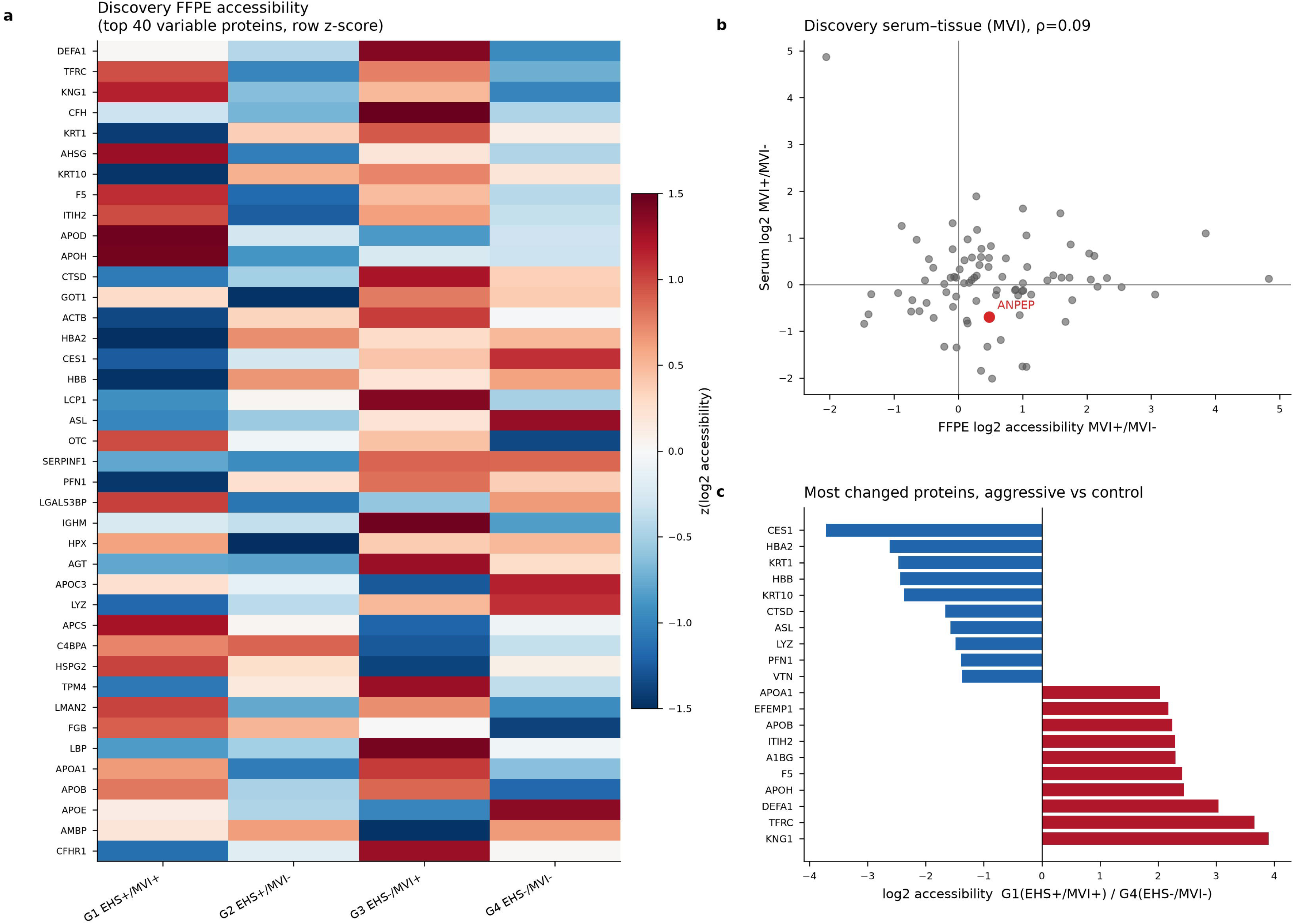
Discovery CPP accessibility profiling across the 2×2 factorial design. (a) Heatmap of tumor (FFPE) accessibility for the 40 most variable common proteins (row z-scored group means) across the four discovery groups (G1, EHS+/MVI+; G2, EHS+/MVI−; G3, EHS−/MVI+; G4, EHS−/MVI−). (b) Serum versus tissue log2 accessibility fold-change for the MVI contrast (MVI+/MVI−) across the 85 common proteins; ANPEP is highlighted (Spearman ρ=0.09). (c) Proteins with the largest accessibility change between the most aggressive group (G1, EHS+/MVI+) and the control group (G4, EHS−/MVI−); blue denotes decreased (closed) and red increased (open) accessibility.

### Macrovascular-invasion–associated accessibility changes propagate from tumor to serum

In the validation cohort, protein-level differential-accessibility testing (two-sided Mann–Whitney U) identified nominal MVI- and EHS-associated changes in both compartments: FGG (decreased) and CTSD, KRT1, LBP (increased) for tumor MVI; ceruloplasmin (increased) and haemoglobin-α and APOE (decreased) for serum MVI; and albumin, HSPA5, lumican, and DEFA1 (all decreased) for serum EHS (**Figure S2**). No protein survived Benjamini–Hochberg correction at this sample size; we therefore report nominal significance together with effect sizes rather than claiming multiplicity-controlled, genome-wide discovery.

The more robust and biologically interpretable signal emerged from translatability—whether a protein’s accessibility moves in the same direction in tumor and serum, the property that ultimately matters for a blood-based test. For the MVI contrast, tumor and serum log2 fold-changes were positively correlated (Spearman ρ=0.21; 59% of proteins concordant in direction; p=0.06), and the concordant proteins were dominated by coagulation and complement factors (FGG, CTSD, LBP, C4BPA) (**Figure 3a**). The EHS contrast showed no such translation (ρ=−0.01; **Figure 3b**), indicating that the vascular-invasion axis, far more than extrahepatic spread, produces protein-complex changes that propagate from the tumor into the circulation. This compartment specificity is consistent with a model in which intravascular—rather than extravascular—tumor dissemination is the dominant driver of the circulating structural signature.

**Figure 3.**
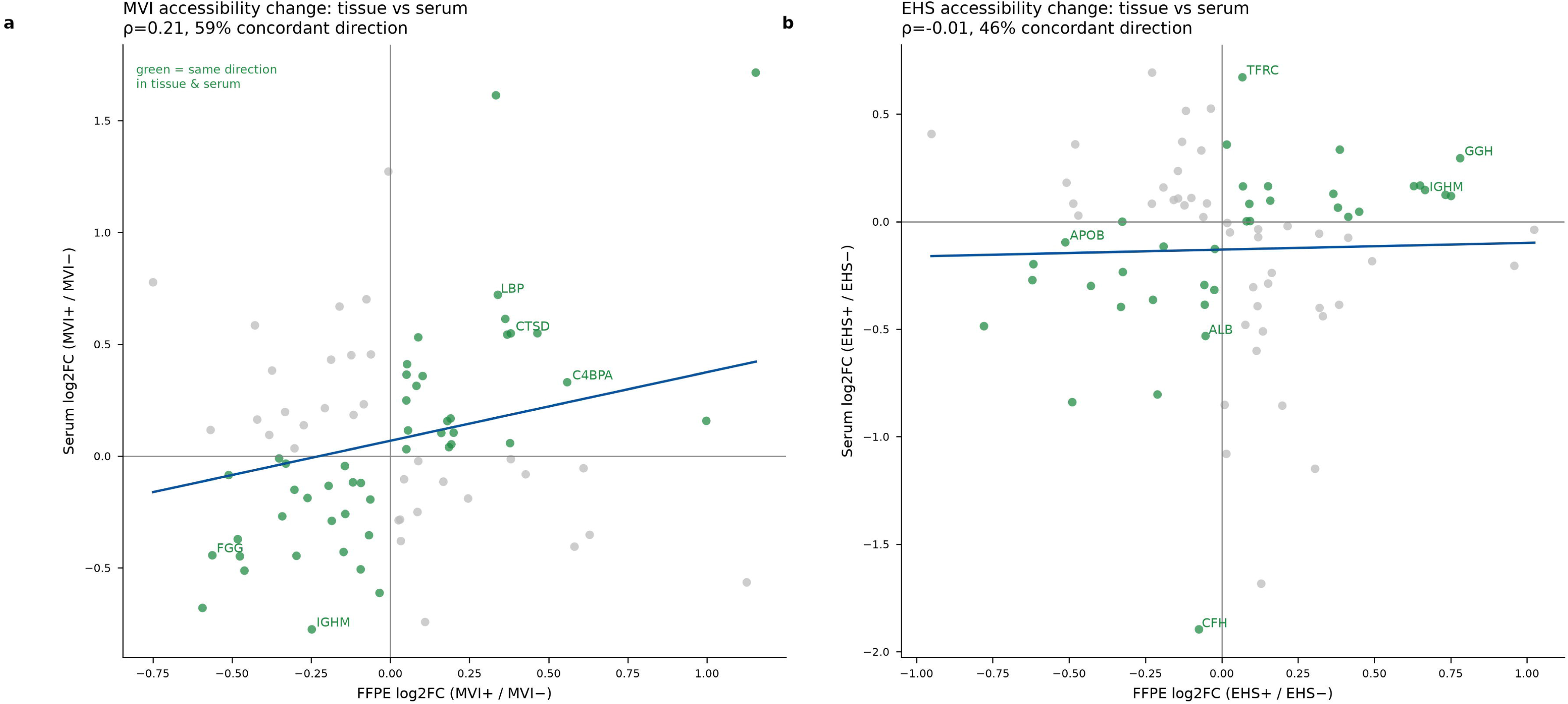
Serum–tissue translatability of accessibility changes distinguishes the MVI and EHS axes. Tumor versus serum log2 accessibility fold-change across the 81-protein panel for the (a) MVI and (b) EHS contrasts. Green points change in the same direction in both compartments; the blue line is the linear fit. The MVI axis shows directional concordance (Spearman ρ=0.21; 59% concordant; p=0.06), driven by coagulation/complement proteins (FGG, CTSD, LBP, C4BPA), whereas the EHS axis does not translate (ρ=−0.01).

### Proteins losing accessibility converge on complement, coagulation, and IGF-transport programs

Proteins losing accessibility in aggressive disease were significantly over-represented in complement and coagulation cascades (KEGG FDR 7.8×10; WikiPathways complement system FDR 3.1×10□³) and, most strongly, in IGF transport and uptake by IGFBPs (FDR 8.7×10□□), as well as scavenger-receptor– mediated ligand uptake, integrin cell-surface interactions, and extracellular-matrix organization (**Figure 4a**). Their STRING physical/functional interaction network formed a densely connected module centered on fibrinogen, serpins, and complement regulators (**Figure 4b**). This is precisely the protein neighborhood expected to be engaged—and therefore rendered less accessible—when intravascular tumor cells activate the platelet, coagulation, and complement systems, providing pathway- and network-level support for the working model.

**Figure 4.**
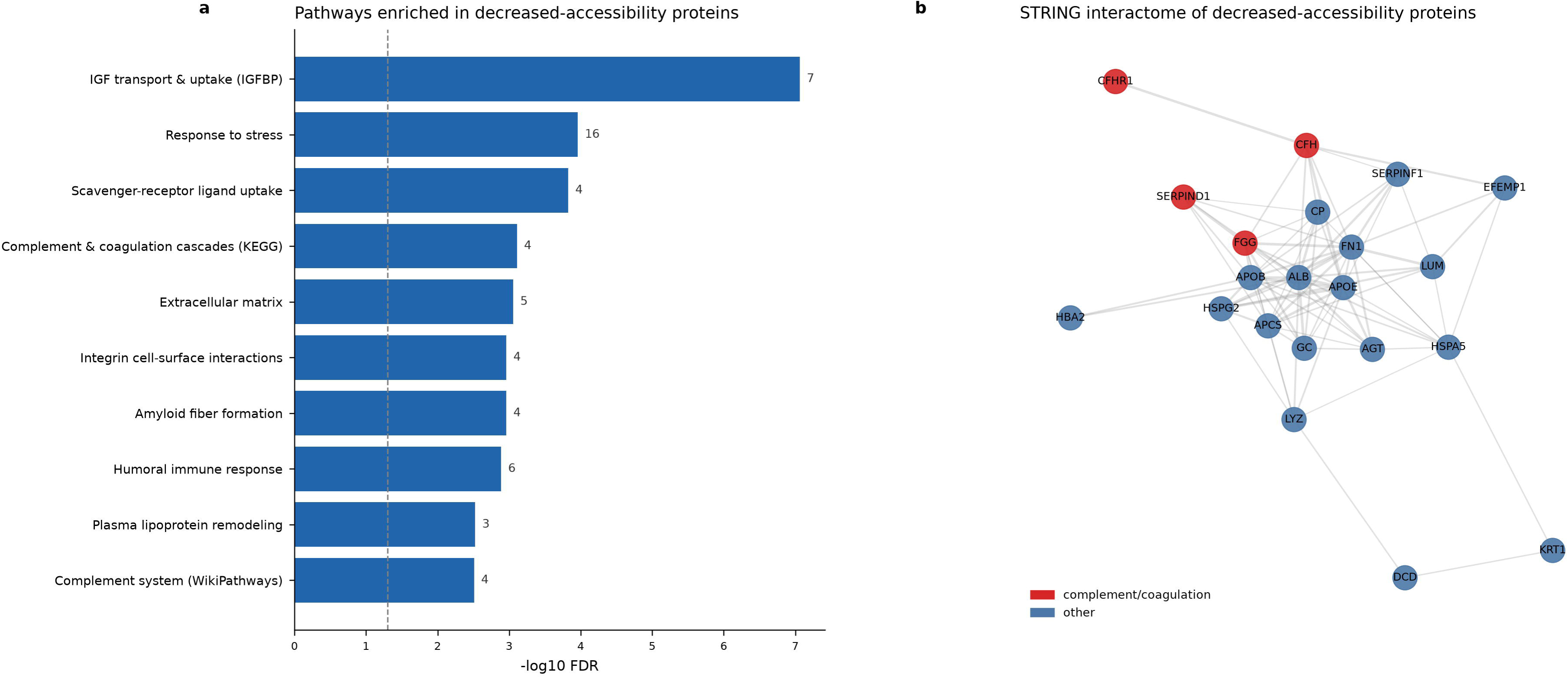
Pathway and network context of decreased-accessibility proteins. (a) Pathways over-represented among proteins with decreased accessibility in aggressive disease (bars, −log10 FDR from STRING functional enrichment; numerals, gene counts; dashed line, FDR=0.05). (b) STRING physical/functional interaction network (minimum combined confidence ≥0.4) of the decreased-accessibility proteins; red nodes denote complement/coagulation members, blue nodes all others.

### ANPEP and coagulation factors report macrovascular invasion through reduced serum accessibility

ANPEP, our prior mechanistic candidate, showed reduced (more closed) serum accessibility across MVI-positive groups relative to the double-negative group, consistent with increased complex engagement of platelet-shed ANPEP in vascular-invasive disease; the difference was directionally supportive but did not reach statistical significance at the available group sizes (**Figure 5a–c**). Serum ANPEP accessibility co-varied modestly with ceruloplasmin accessibility (ρ=0.35, p=0.09), linking it to the broader acute-phase/complement module. The statistically stronger MVI-associated serum markers were coagulation and inflammation proteins—fibrinogen-γ, cathepsin D, and LPS-binding protein (**Figure 5d–f**)— reinforcing a coagulation-centred model in which the circulating accessibility signature is anchored by haemostatic and complement components.

**Figure 5.**
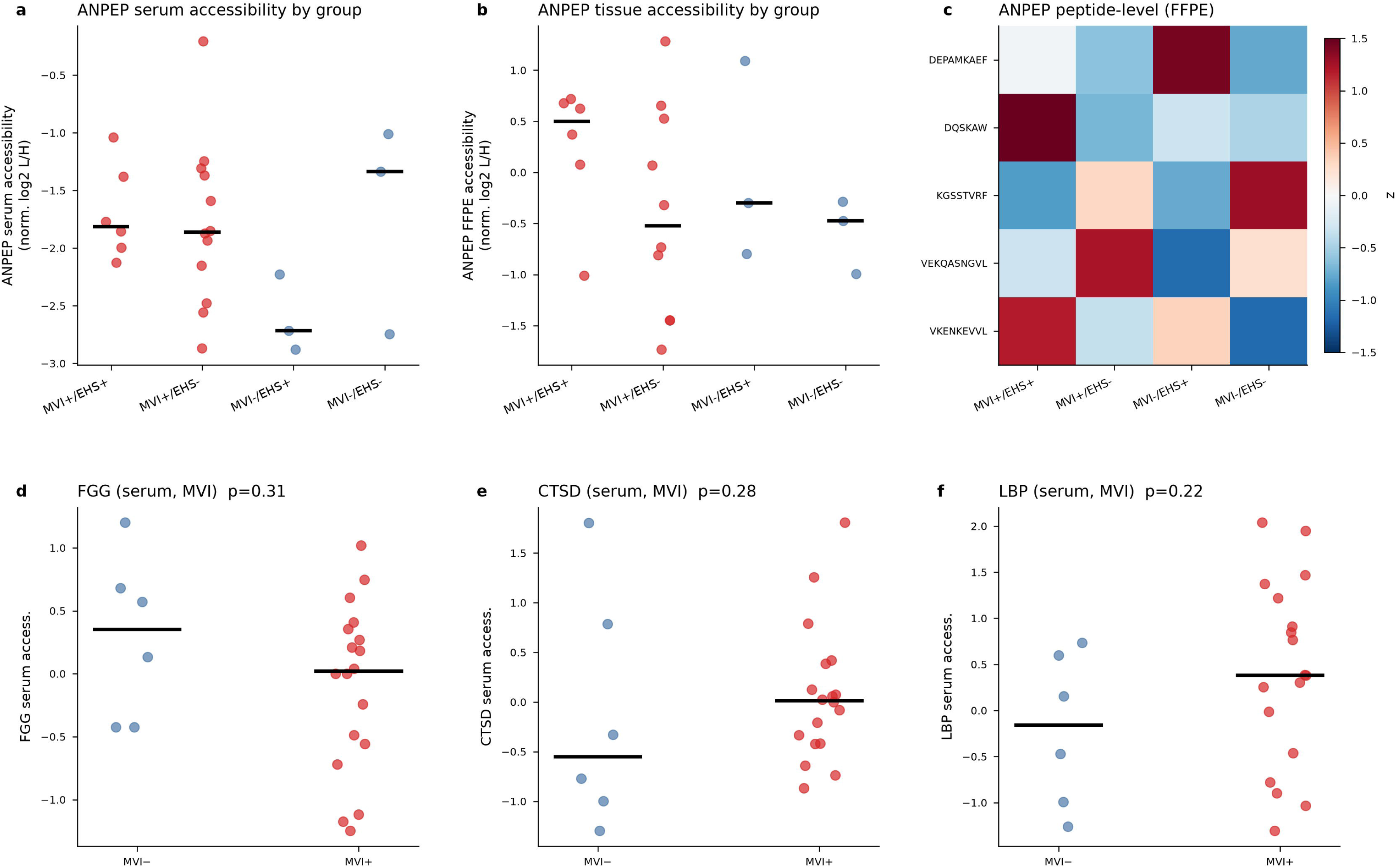
ANPEP and leading MVI-associated markers show reduced serum accessibility. ANPEP accessibility by group in (a) serum and (b) tumor (horizontal bars, group medians). (c) ANPEP peptide-level accessibility (FFPE group means, z-scored). Serum accessibility by MVI status for the leading coagulation/inflammation markers (d) FGG, (e) CTSD, and (f) LBP; horizontal bars are medians and p values are from two-sided Mann–Whitney U.

### A six-protein serum accessibility signature discriminates macrovascular invasion and stratifies survival

A parsimonious six-protein serum accessibility classifier (CP, HBA2, APOE, FGG, LBP, CTSD) discriminated macrovascular invasion with a leave-one-out cross-validated AUC of 0.80; the single strongest markers were ceruloplasmin (AUC 0.83) and haemoglobin-α (0.77) accessibility (**Figure 6a–c**; **Table 2**). An analogous EHS classifier performed considerably less well (AUC 0.61), mirroring the compartment-translatability result and again distinguishing the MVI from the EHS axis. Clinically, macrovascular invasion trended toward shorter overall survival (log-rank p=0.06), and accessibility-derived scores showed concordant but non-significant separation at this cohort size (**Figure 6d–f**). Together, these results establish proof-of-concept that structural, accessibility-based serum proteomics carries MVI-relevant information that is both biologically coherent and, in cross-validation, discriminative.

**Figure 6.**
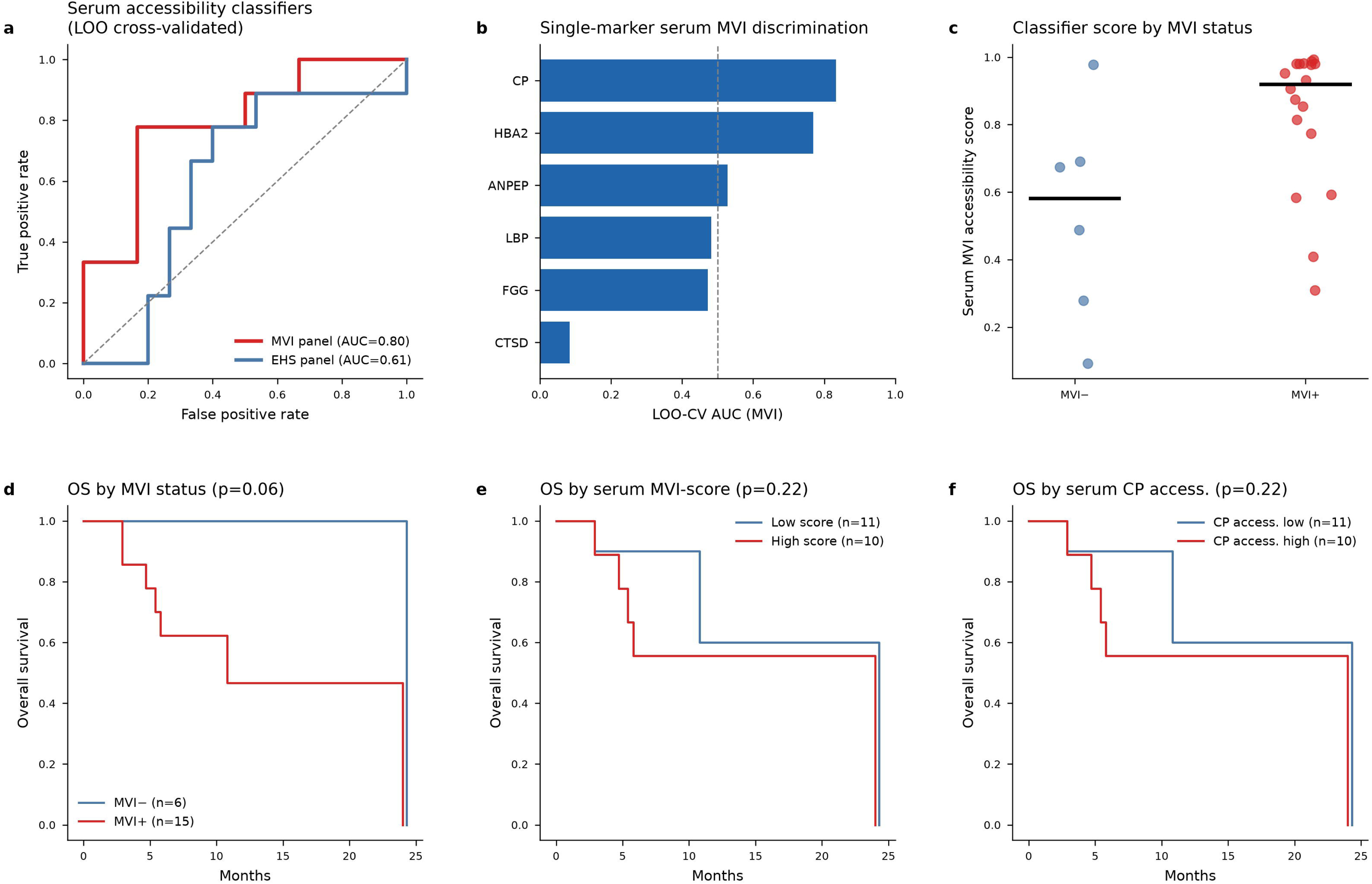
Serum accessibility biomarkers discriminate MVI and stratify survival. (a) Leave-one-out cross-validated ROC curves for the serum MVI (6-marker) and EHS (5-marker) accessibility classifiers. (b) Single-marker LOO-CV AUC for MVI discrimination. (c) Serum MVI accessibility score by MVI status (horizontal bars, group medians). Kaplan–Meier overall survival stratified by (d) MVI status, (e) serum MVI accessibility score (median split), and (f) serum ceruloplasmin (CP) accessibility (median split); p values are from the log-rank test.

**Table 2.** Serum accessibility biomarker performance for macrovascular invasion. Leave-one-out cross-validated (LOO-CV) discrimination of MVI status from serum protein accessibility. Multi-marker panels are L2-regularized logistic-regression classifiers; single-marker AUCs are shown for the leading proteins.

| Protein | UniProt | Serum<br>MVI<br>log2FC | Serum<br>MVI p | Direction | LOO-CV<br>AUC<br>(MVI) |
| --- | --- | --- | --- | --- | --- |
| CP | P00450 | 1.270 | 0.001 | ↑open | 0.830 |
| HBA2 | P69905 | -0.740 | 0.012 | ↓closed | 0.770 |
| ANPEP | P15144 | 0.550 | 0.224 | ↑open | 0.530 |
| LBP | P18428 | 0.720 | 0.224 | ↑open | 0.480 |
| FGG | P02679-2 | -0.440 | 0.310 | ↓closed | 0.470 |
| CTSD | P07339 | 0.550 | 0.280 | ↑open | 0.080 |
| 6-marker MVI panel | CP, HBA2, APOE, FGG, LBP, CTSD |  |  |  | 0.800 |
| 5-marker EHS panel | ALB, HPX, APOB, GC, IGHM |  |  |  | 0.610 |

*AUC, area under the receiver-operating-characteristic curve; EHS, extrahepatic spread; LOO-CV, leave-one-out cross-validation; MVI, macrovascular invasion.*

## Discussion

We show that a structural-proteomics readout—protein accessibility measured by covalent proteome painting—captures features of aggressive HCC that are, by construction, invisible to abundance-based assays^19^. The central positive finding is directional: macrovascular invasion is accompanied by decreased accessibility of coagulation and complement proteins, and these changes are shared between tumor tissue and serum^24^. This compartment translatability is the property that matters for a blood test, and it is specific to the MVI axis; extrahepatic spread, despite its clinical aggressiveness, did not produce a comparably translatable structural signature^25^.

The biology fits a coherent model. Intravascular tumor dissemination activates platelets and the coagulation and complement cascades, promoting assembly of protein complexes—fibrinogen networks, complement-regulator assemblies, and serpin–protease pairs—in which the participating proteins bury the very lysine and N-terminal surfaces that CPP labels. The result is a coordinated loss of accessibility across an interacting module, exactly the pattern we observe by enrichment and network analysis. ANPEP, shed from activated platelet membranes, is a natural member of this program, and its reduced serum accessibility in MVI-positive disease, although not individually significant here, is consistent with the hypothesis that motivated the study.

Several limitations bound these conclusions. The discovery phase used two patients per group, and the validation cohort of 22 tumors and 22 sera, while adequate to demonstrate translatability and cross-validated classification, is underpowered for genome-wide, multiplicity-controlled discovery; no single protein survived FDR correction, and we have been explicit in reporting nominal significance with effect sizes. The cohort is advanced-stage and HBV-predominant, so generalization to earlier disease or other etiologies remains untested. Accessibility ratios can in principle be influenced by protein abundance and by depletion efficiency; although the light/heavy SIS design and per-sample normalization mitigate this, orthogonal validation of specific complexes—for example by co-immunoprecipitation or cross-linking MS—would strengthen the mechanistic claims. Finally, survival associations are hypothesis-generating at this event count.

Despite these constraints, the study makes a methodological and biological point that we believe is broadly relevant: interaction-state proteomics can reveal disease-associated protein-complex remodeling that abundance misses, and at least for macrovascular invasion in HCC, that remodeling reaches the blood. A coagulation/complement-anchored serum accessibility signature—led by ceruloplasmin, fibrinogen-γ, and related factors—is a concrete, testable candidate for non-invasive detection of vascular-invasive HCC^26^, and the CPP-to-MRM workflow demonstrated here provides a scalable route to validate it in larger, prospectively collected cohorts.

## Materials and Methods

### Study design, participants, and ethical approval

This two-stage study comprised an unbiased discovery phase and a targeted validation phase. Treatment-naïve patients with histologically or radiologically confirmed HCC were enrolled, and matched formalin-fixed paraffin-embedded (FFPE) tumor tissue and pre-treatment serum were obtained from each participant. Macrovascular invasion (MVI) and extrahepatic spread (EHS) were determined from integrated clinical, radiologic, and pathologic staging according to standard criteria. For discovery, eight patients were selected in a balanced 2×2 factorial design (MVI± × EHS±, two patients per group). For validation, all cross-compartment comparisons were harmonized to the same 22 patients with matched FFPE tumor and serum. The study was conducted in accordance with the Declaration of Helsinki and approved by the Institutional Review Board of Seoul National University Hospital (IRB approval no. 2211-152-1381); written informed consent was obtained from all participants, and all clinical data were de-identified prior to analysis.

### Serum depletion and covalent proteome painting (discovery)

Protein accessibility was labeled by reductive dimethylation, which covalently modifies solvent-accessible lysine ε-amines and protein N-termini while sparing residues protected within folded structures or at protein–protein interfaces; labeling intensity thereby reflects local accessibility, with high signal indicating open binding sites and low signal indicating closed, complex-engaged sites. Prior to labeling, serum was depleted of the 14 most-abundant plasma proteins using a High-Select Top14 Abundant Protein Depletion resin to expose the lower-abundance interactome. Labeled proteins were digested and the resulting peptides analyzed by data-dependent LC–MS/MS on a Q Exactive hybrid quadrupole-Orbitrap mass spectrometer operated at a resolution of 120,000 with mass accuracy <3 ppm. Protein-level intensities were quantified by the MSFragger program^27^. Proteins detected in both the tumor and serum compartments defined the common proteome from which the targeted panel was constructed.

### Targeted MRM assay and accessibility quantification (validation)

The 81-protein panel was quantified by multiple reaction monitoring-mass spectrometry (MRM-MS)^28-31^. For each targeted peptide, an isotopically heavy stable-isotope-standard (SIS) peptide was spiked as an internal reference, and both the endogenous light and the heavy transitions were monitored (296 peptides; 3,717 paired light/heavy transitions). Accessibility for each transition was defined as the light/heavy intensity ratio; peptide-level accessibility was computed as the summed light over the summed heavy fragment-ion intensities, and protein-level accessibility as the median across a protein’s constituent peptides. Protein accessibility values were log2-transformed and median-centered per sample to remove global labeling and loading variation prior to statistical comparison. Assay performance was assessed by transition-level coefficient of variation, heavy-versus-light intensity correlation, sample–sample correlation, and per-sample panel completeness (**Figure S1**).

### Differential accessibility and serum–tissue translatability

Differential accessibility between MVI or EHS strata was tested by two-sided Mann–Whitney U on log2 accessibility, with Benjamini–Hochberg control of the false-discovery rate across the 81 proteins. Effect sizes are reported as log2 fold-change (median difference) and rank-biserial correlation. Serum–tissue translatability was assessed, for each contrast, as the Spearman correlation between tumor and serum contrast log2 fold-changes together with the fraction of proteins concordant in the direction of change.

### Pathway enrichment and interaction-network analysis

Functional enrichment of the proteins with decreased accessibility in aggressive disease was performed using the STRING database functional-enrichment interface, querying KEGG, Reactome, WikiPathways, and Gene Ontology annotation sets; enrichment significance is reported as the Benjamini–Hochberg false-discovery rate^32^. Physical and functional interactions among the same proteins were retrieved from STRING at a minimum combined confidence of 0.4 and visualized as an interaction network, with complement and coagulation members highlighted.

### Biomarker classification and survival analysis

Serum accessibility classifiers for MVI and EHS were built as L2-regularized logistic-regression models on standardized accessibility features and evaluated by leave-one-out cross-validation, with the area under the receiver-operating-characteristic curve (AUC) computed on out-of-fold predictions to avoid optimistic bias; single-marker AUCs were computed analogously. Overall survival was estimated by the Kaplan– Meier method and compared between strata by the log-rank test; continuous accessibility scores were dichotomized at the cohort median.

### Statistical analysis and reproducibility

Clinical continuous variables are summarized as median (interquartile range) and compared by Mann– Whitney U; categorical variables are compared by Fisher’s exact test. All analyses were performed in Python using scipy, statsmodels, scikit-learn, and lifelines. Unless otherwise stated, tests were two-sided and nominal significance was defined at p<0.05, with multiplicity explicitly noted where controlled.

## Data and code availability

The mass spectrometry data have been deposited to Panorama Public (https://panoramaweb.org/), and are accessible at https://panoramaweb.org/vHmjuf.url. All analysis scripts used in this study are publicly available on GitHub (https://github.com/kimlab-cnu/HCCstructMarker). All quantitative accessibility matrices, per-contrast differential statistics, pathway-enrichment results, biomarker-performance metrics, and de-identified clinical data are provided in the accompanying Supplementary Data (**Supplementary Tables S1–S7**). Additional data are available from the corresponding author on reasonable request.

## Acknowledgement

We acknowledge the use of Claude Opus 4.8 solely for linguistic refinement and grammatical corrections in manuscript preparation. All scientific content, data analysis, and intellectual contributions presented herein were developed independently by the authors without the use of generative AI tools.

## Author Contributions

A.S., M.H., S.J.Y., and H.K.: conceptualization, methodology, experimental analysis, data analysis, visualization; E.J.C., J.J., E.H., Y.C., J.P, H.L., and S.P: data analysis; S.J.Y., and H.K.: writing-original draft, conceptualization, project administration, resources, supervision, writing-review & editing. All authors have read and approved the final manuscript.

---

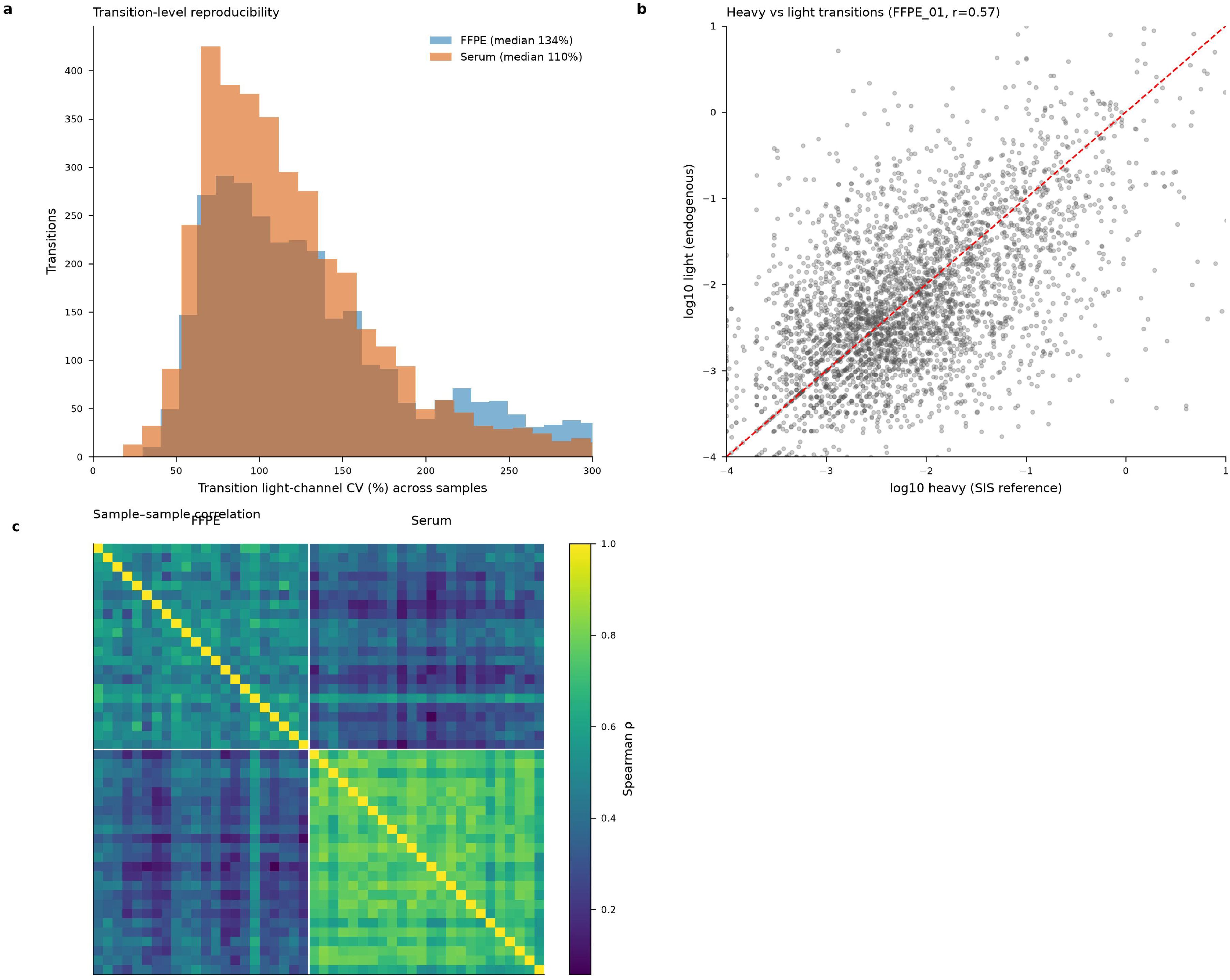

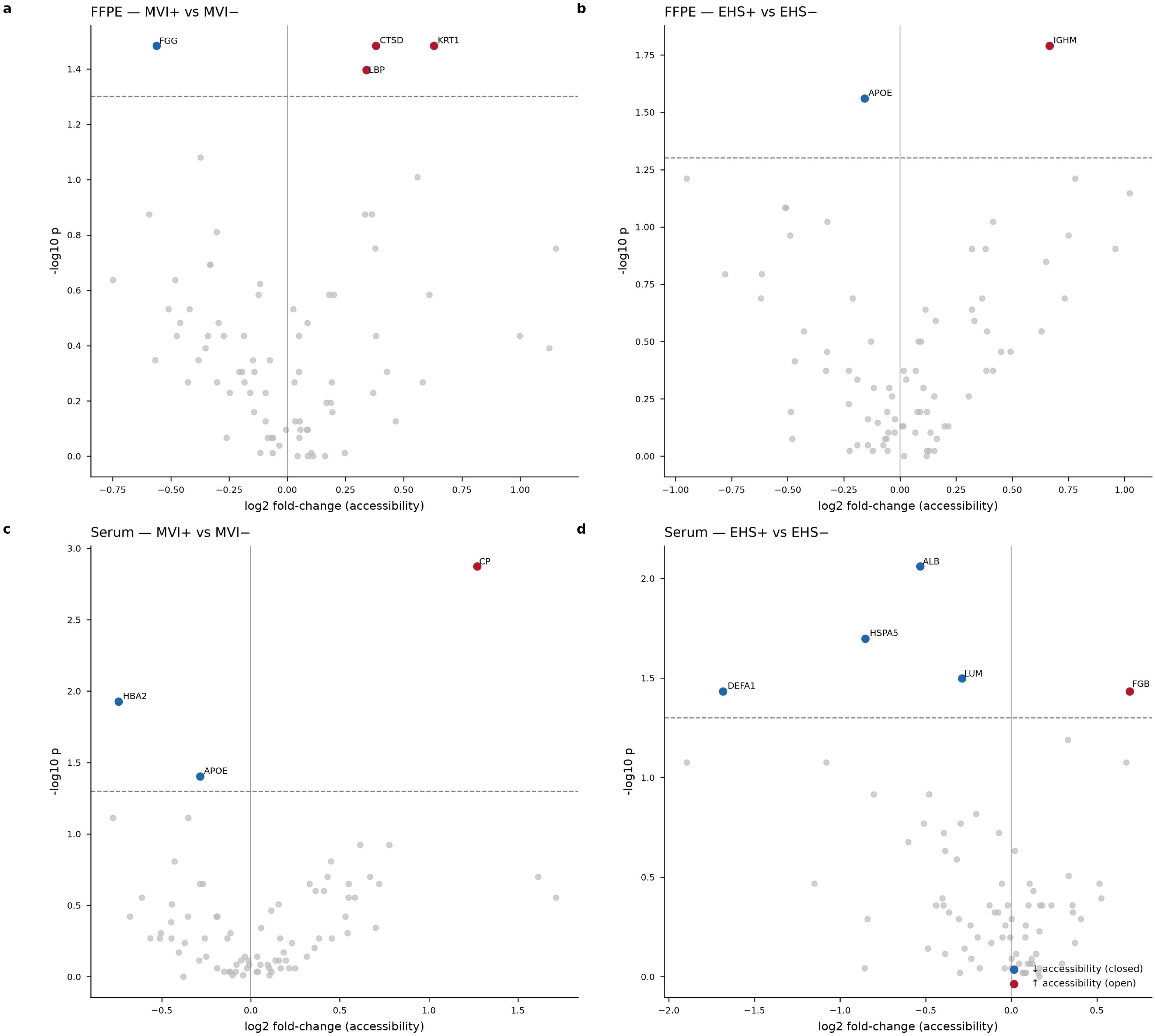

