## Supplemental Information for "Structural proteomics reveals a coagulation–complement accessibility signature of macrovascular invasion in hepatocellular carcinoma"

**Supplementary Figures**

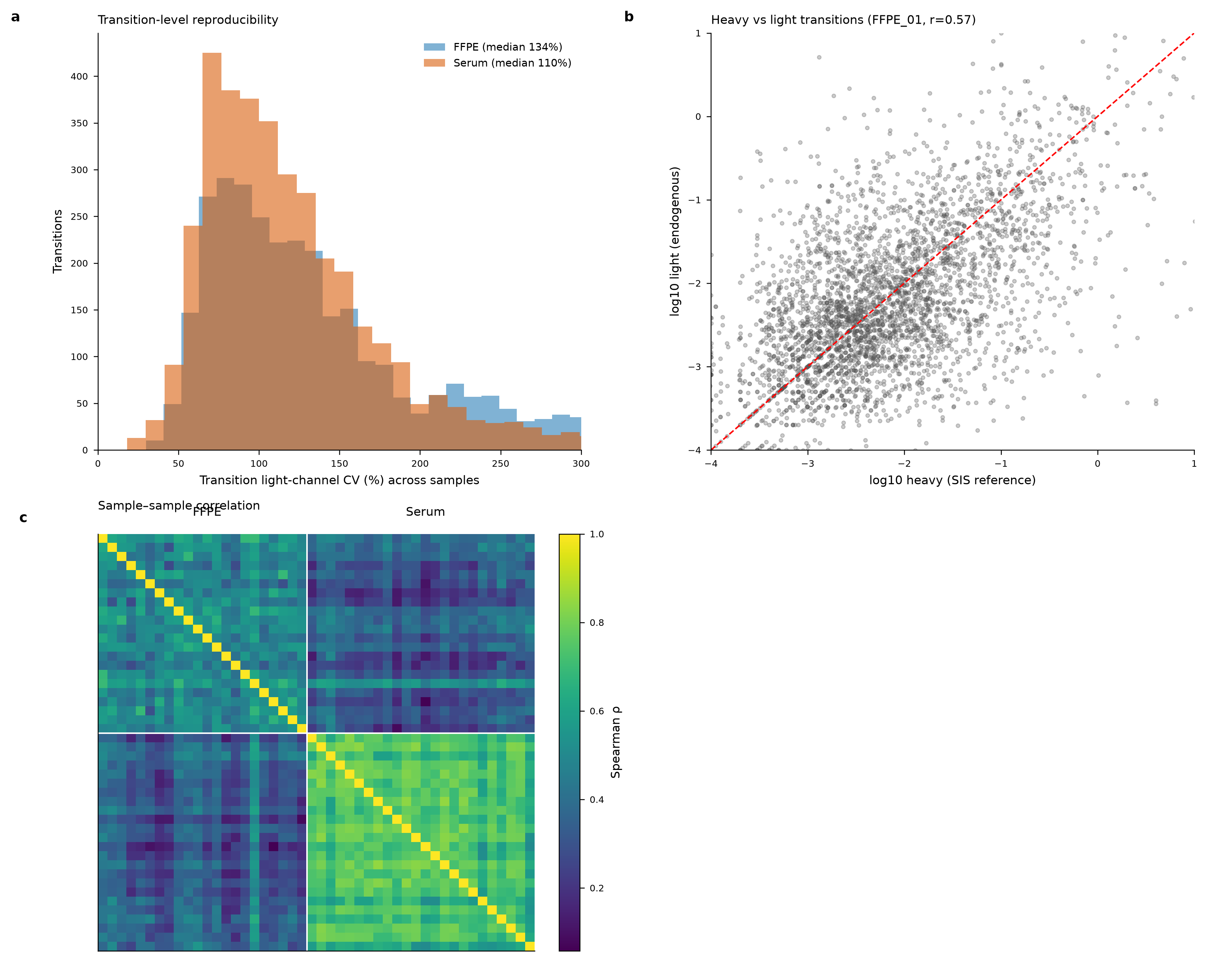

**Figure S1. Assay quality control.** (a) Transition-level light-channel CV across samples (FFPE and serum). (b) Heavy (SIS) versus light (endogenous) transition intensities for a representative sample. (c) Sample–sample Spearman correlation.

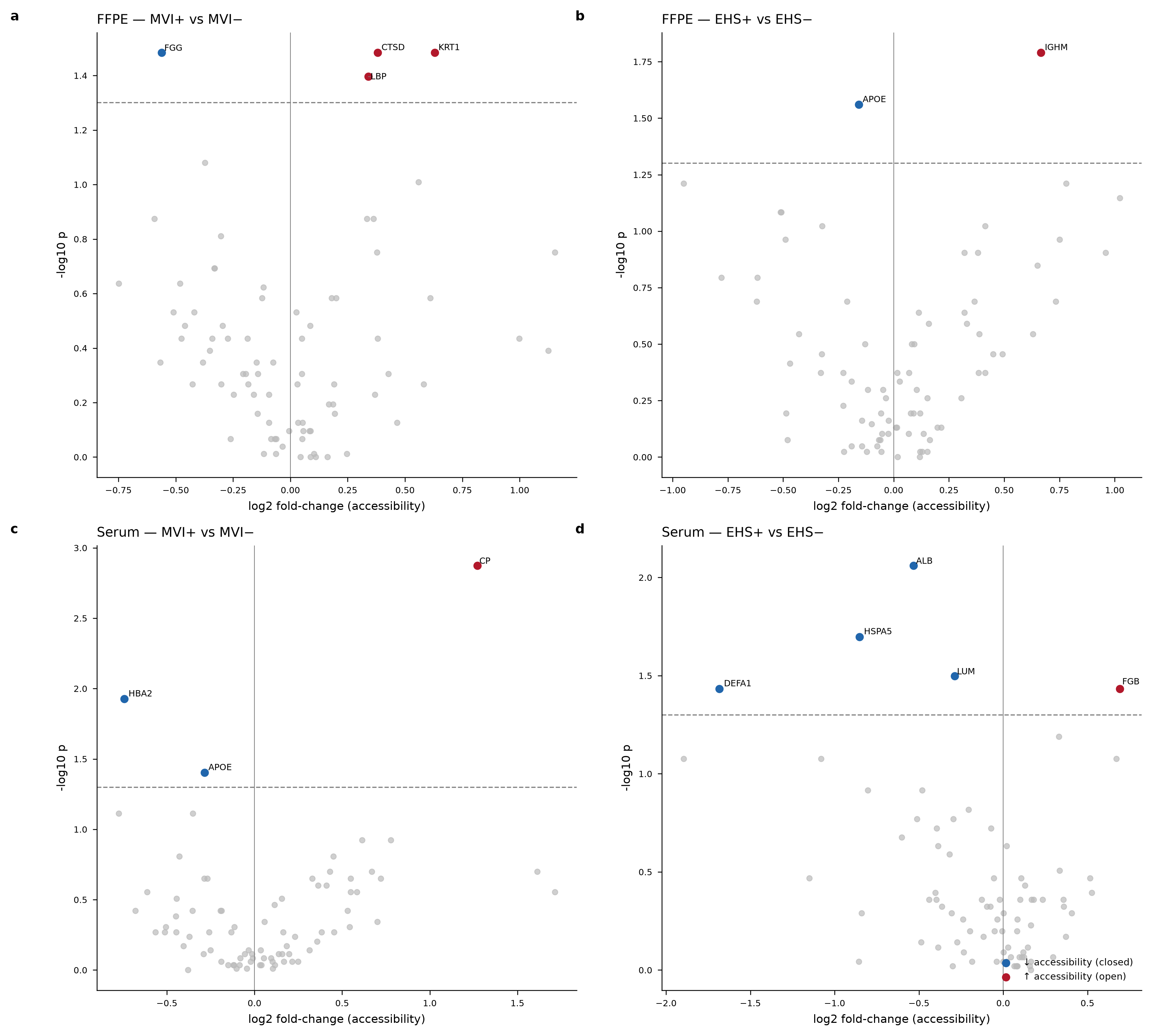

**Figure S2. Protein-level differential accessibility in the validation cohort.**

Volcano plots of protein-level log2 fold-change versus −log10 p (two-sided Mann–Whitney U) for (a) FFPE MVI+ vs MVI−, (b) FFPE EHS+ vs EHS−, (c) serum MVI+ vs MVI−, and (d) serum EHS+ vs EHS−. Proteins at nominal p<0.05 are colored (blue, decreased/closed accessibility; red, increased/open accessibility); the dashed line marks p=0.05. No protein survived Benjamini–Hochberg FDR<0.05 at this sample size, so nominal signals are reported together with effect sizes.
